# Evolution of populations with strategy-dependent time delays

**DOI:** 10.1101/865071

**Authors:** Jacek Miȩkisz, Marek Bodnar

## Abstract

We address the issue of stability of coexistence of two strategies with respect to time delays in evolving populations. It is well known that time delays may cause oscillations. Here we report a novel behavior. We show that a microscopic model of evolutionary games with a unique mixed evolutionarily stable strategy (a globally asymptotically stable interior stationary state in the standard replicator dynamics) and with strategy-dependent time delays leads to a new type of replicator dynamics. It describes the time evolution of fractions of the population playing given strategies and the size of the population. Unlike in all previous models, an interior stationary state of such dynamics depends continuously on time delays and at some point it might disappear, no cycles are present. In particular, this means that an arbitrarily small time delay changes an interior stationary state. Moreover, at certain time delays, there may appear another interior stationary state.

**Author summary:** Social and biological processes are usually described by ordinary or partial differential equations, or by Markov processes if we take into account stochastic perturbations. However, interactions between individuals, players or molecules, naturally take time. Results of biological interactions between individuals may appear in the future, and in social models, individuals or players may act, that is choose appropriate strategies, on the basis of the information concerning events in the past. It is natural therefore to introduce time delays into evolutionary game models. It was usually observed, and expected, that small time delays do not change the behavior of the system and large time delays may cause oscillations. Here we report a novel behavior. We show that microscopic models of evolutionary games with strategy-dependent time delays, in which payoffs appear some time after interactions of individuals, lead to a new type of replicator dynamics. Unlike in all previous models, interior stationary states of such dynamics depend continuously on time delays. This shows that effects of time delays are much more complex than it was previously thought.

## Introduction

Many social and biological processes can be modeled as systems of interacting individuals within the framework of evolutionary game theory [9,11,15–17,22,23,26,29]. The evolution of very large (infinite) populations can be then given by differential replicator equations which describe time changes of fractions of populations playing different strategies [10,11,25,26]. It is usually assumed (as in the replicator dynamics) that interactions between individuals take place instantaneously and their effects are immediate. In reality, all social and biological processes take a certain amount of time. Results of biological interactions between individuals may appear in the future, and in social models, individuals or players may act, that is choose appropriate strategies, on the basis of the information concerning events in the past. It is natural therefore to introduce time delays into evolutionary game models.

It is well known that time delays may cause oscillations in dynamical systems [2–5]. One usually expects that interior equilibria of evolving populations, describing coexisting strategies or behaviors, are asymptotically stable for small time delays and above a critical time delay, where the Hopf bifurcation appears, they become unstable, evolutionary dynamics exhibits oscillations and cycles. Here we report a novel behavior - continuous dependence of equilibria on time delays.

Effects of time delays in replicator dynamics were discussed in [6–8,12,13,18–21,24,27,28] for games with an interior stable equilibrium (an evolutionarily stable strategy [15,16]). In [24], the authors discussed the model, where individuals at time t imitate a strategy with a higher average payoff at time *t* – *τ* for some time delay *τ*. They showed that the interior stationary state of the resulting time-delayed differential equation is locally asymptotically stable for small time delays and for big ones it becomes unstable, there appear oscillations. In [6], we constructed a different type of a model, where individuals are born *τ* units of time after their parents played. Such a model leads to system of equations for the frequency of the first strategy and the size of the population. We showed the absence of oscillations - the original stationary point is globally asymptotically stable for any time delay.

Here we modify the above model by allowing time delays to depend on strategies played by individuals.

Recently there were studied models with strategy-dependent time delays. In particular, Moreira et al. [20] discussed multi-player Stag Hunt game with time delays, Ben Khalifa et al. [8] investigated asymmetric games in interacting communities, Wesson and Rand [27] studied Hopf bifurcations in two-strategy delayed replicator dynamics.

Here we discuss effects of strategy-dependent time delays on the long-run behavior of a biological-type model of certain two-player games with two strategies. We report a novel behavior. We showed that an arbitrarily small time delay may change an interior stationary state. Moreover, at certain time delays, an interior stationary state may disappear and the pure strategy becomes globally asymptotically stable or there may appear another interior stationary state. In general, the effect of strategy-dependent time delays might be summarized as follows: bigger a time delay of a given strategy, smaller its proportion in an asymptotically stable stationary state.

## Materials and methods

### Replicator dynamics

We assume that our populations are haploid, that is the offspring have identical phenotypic strategies as their parents. We consider symmetric two-player games with two strategies, A and B, given by the following payoff matrix:

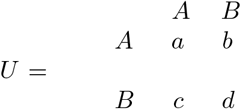

where the *ij* entry, *i, j* = *A, B*, is the payoff of the first (row) player when it plays the strategy *i* and the second (column) player plays the strategy *j*. We assume that both players are the same and hence payoffs of the column player are given by the matrix transposed to *U*; such games are called symmetric.

Let us assume that during a time interval of the length *ε*, only an *ε*-fraction of the population takes part in pairwise competitions, that is plays games. Let *p_i_*(*t*), *i* = *A, B*, be the number of individuals playing at the time t the strategy *A* and *B* respectively, *p*(*t*) = *p_A_*(*t*) + *p_B_* (*t*) the total number of players and 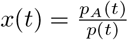 the fraction of the population playing *A*. Let

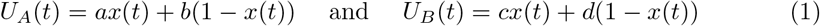

be average payoffs of individuals playing A and B respectively.

We assume *d < b < a < c*, so there exists a unique mixed evolutionarily stable strategy ([15,16]), 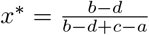, which represents the equilibrium fraction of an infinite population playing *A* [11,26]. In the replicator dynamics, *x*\* is globally asymptotically stable and there are two unstable stationary states: *x* = 0 and *x* = 1.

Now we would like to take into account that individuals are born some units of time after their parents played. We assume that time delays depend on strategies and are equal to *t* – *τ_A_* or *t* – *τ_B_* respectively.

We propose the following equations:

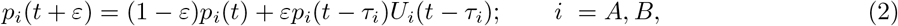

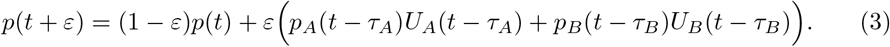

We divide (2) by (3) for *i* = *A*, obtain the equation for *x*(*t* + *ε*) ≡ *x_A_*(*t* + *ε*, subtract *x*(*t*), divide the difference by *ε*, take the limit *ε* → 0, and get an equation for the frequency of the first strategy,

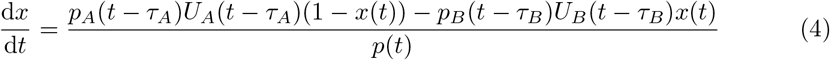

which can be also written as

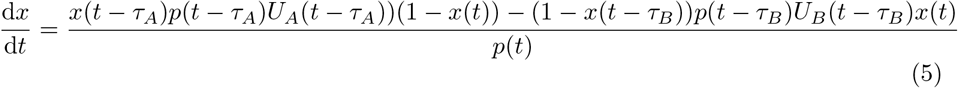

Let us notice that unlike in the standard replicator dynamics, the above equation for the frequency of the first strategy is not closed, there appear in it a variable describing the size of the population at various times. One needs corresponding equations for the population size. From (2) and (3) we have

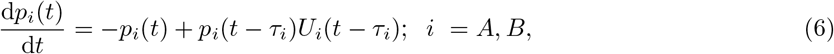

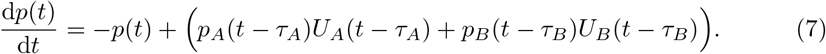

To trace the evolution of the population, we have to solve the system of equations, (5,7), together with initial condition on the interval [–*τ_M_*, 0], where *τ_M_* = max{*τ_A_,τ_B_*}. We assume that

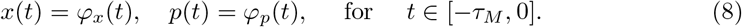

It can be shown (see Proposition 2 in SI) that if the initial functions *φ_x_, φ_p_* are continuous on [*τ_M_*, 0) and non-negative, then there exists a unique, non-negative solution of the system ((5), (7)) with the initial conditions (8) which is well defined on the interval [0, +∞).

### Stationary state

We derive here an equation for a stationary state of the frequency of the first strategy. Let us assume that there exists a stationary frequency 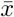 such that 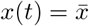 for all *t* ≥ 0 and for some suitably chosen function *p*(*t*). Then average payoffs of each strategy are constant and are equal to

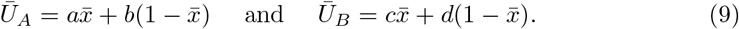

Thus, equation (7) becomes a linear delay differential equation,

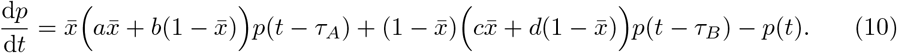

Note, that solutions of (10) with non-negative initial conditions are non-negative. This implies that the leading eigenvalue of this equation is real. The eigenvalues λ of (10) satisfy

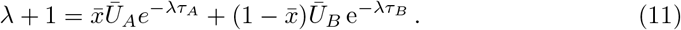

Assume now that λ is a solution of (11) (of course λ depends on 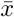) and *p*(*t*) = *p*_0_ exp(λ*t*) for some *p*_0_.

We plug such *p* into (4) and we get two stationary solutions of (4) (i.e. such that the right-hand side of (4) is equal to 0), 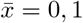 and possibly interior ones — solutions to

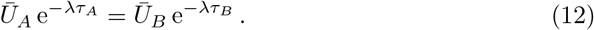

If *τ_A_* = *τ_B_*, then (12) gives us a mixed Nash equilibrium (an evolutionarily stable strategy) of the non-delayed replicator dynamics,

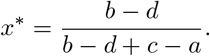

If *τ_A_* ≠ *τ_B_*, we have then 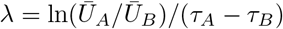. Plugging it into (11) we conclude that 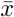 satisfies an equation 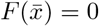, where

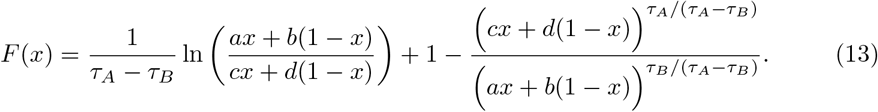

The above reasoning leads immediately to the following proposition which explains in what sense 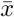 is a stationary state of the replicator dynamics ((5), (7)).

#### Proposition 1.

*Assume that τ_A_ ≠ τ_B_. Let 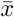 be a solution to F*(*x*) = 0, *where F is defined by* (13) *and let*

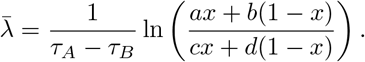

*Then the functions*

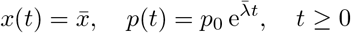

*are solutions of the system (*(5)–(7)*) with the initial functions*

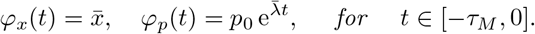

## Results

To exhibit new types of behavior of evolutionary dynamics with strategy-dependent time delays, we will discuss now two particular games.

### Example 1

Here we consider a game with the following payoff matrix,

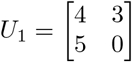

We have that *x*\* = 0.75 is the stable stationary state of the replicator dynamics.

When delays are parametrized in the following form, *τ_A_* = *τ, τ_B_* = 2*τ* and *τ* increases, then the interior stationary state increases until it disappears at some value of *τ*. Above that point, *x* = 1 becomes globally asymptotically stable (see Fig. 1A). In the case of *τ_A_* = 2*τ, τ_B_* = *τ*, the interior stationary state decreases and approaches the value 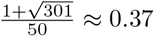 (see Fig. 1C).

We see in Fig. 1B, that for fixed *τ_B_* = 2, there is a value of *τ_A_* below which there is no interior stationary point; *x* = 1 is globally asymptotically stable. Above this point, the interior stationary state decreases.

In Fig 1D we present the interior stationary state as function of *τ_A_* and *τ_B_*. If *τ_A_* = *τ_B_*, then the stable stationary frequency does not depends on time delay and is equal to *x*\* = 0.75. For fixed *τ_B_*, if *τ_A_* ≫ *τ_B_*, the stationary state decreases approaching the value 0.2 (that can be easily calculated taking a limit *τ_A_* → + ∞ in (13)). Similarly, analyzing the graph of the function *F* for *τ_B_* =0 one can easily find that the stationary state decreases in this case from 0.75 for *τ_A_* = 0 to 0.2 for *τ_A_* → +∞. On the other hand, if we fixed *τ_B_* then the stationary state is a decreasing function of *τ_A_* (see Fig. 1B). Additionally, if *τ_B_* is large enough (greater that ln(5/4)/3, see Proposition 9 in SI), then there exists a value 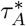 of *τ_A_*, such that for 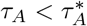 there exists no interior stable frequency and all players are going to play the first strategy. The critical value 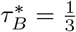 ln 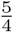 for *τ_A_* = 0 depends on *τ_A_* almost linearly (see the implicit formula (18) in the SI for the relation between 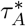, and 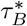).

**Fig 1.**
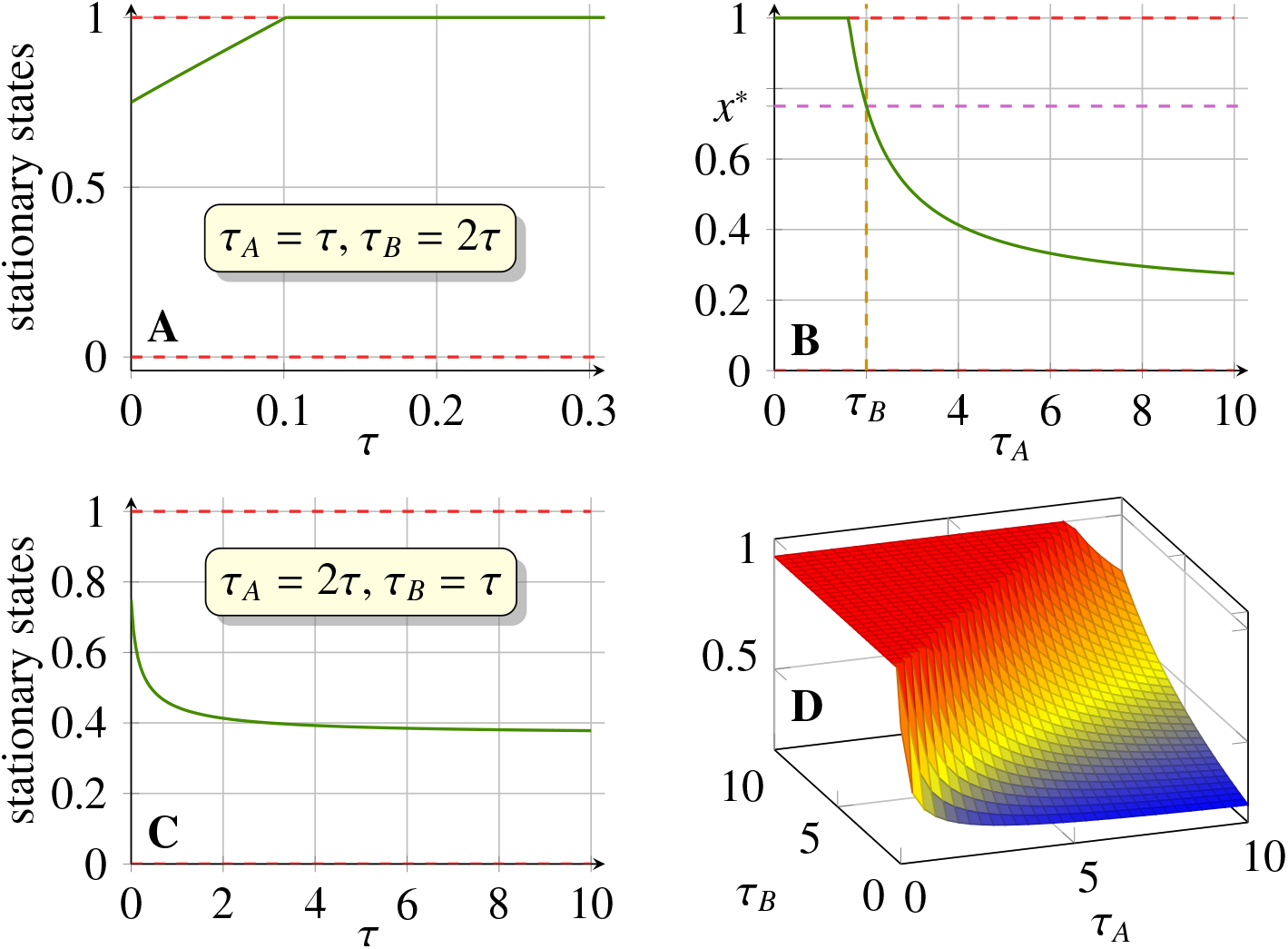
Numerical solutions of *F*(*x*) = 0 for the matrix *U*_1_. Panel **A**: interior stationary state as a function of *τ*, when *τ_A_* = 2*τ, τ_B_* = *τ*. Panel **B**: interior stationary state as a function of *τ_A_*, while *τ_B_* = 2 is fixed. Panel **C**: interior stationary state as a function of *τ*, when *τ_A_* = *τ*, *τ_B_* = 2*τ*. Panel **D**: interior stationary state as a function of *τ_A_* and *τ_B_*.

### Example 2

Here we study a game with the following payoff matrix,

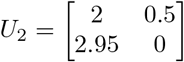

Now, *x*\* ≈ 0.345 is the stable stationary state of the replicator dynamics.

In Fig 2A we set *τ_A_* = *τ, τ_B_* = 2*τ*. We see that here there exists a threshold *τ*\* ≈ 1.1 such that for *τ* < *τ*\* there exists a unique interior stationary state. Numerical simulations suggest that this state is stable. For *τ* > *τ*\* there are two interior stationary states. Numerical simulations suggest that if an initial frequency of cooperation strategy is large enough, then the detection strategy is eliminated.

In Fig. 2B we present the interior stationary state as a function of *τ, τ_A_* = 2*τ*, *τ_B_* = *τ*. In this case, the stationary state is almost constant. In fact, for this set of parameters, one can easily see that the function F is a decreasing one as both terms of F decrease. Because for chosen *τ_A_* and *τ_B_*, only the first term depends on *τ* (and it decreases with increasing *τ*) one can deduce that the stationary state is a decreasing function of *τ*. For *τ* → +∞ it converges to the positive solution of the quadratic equation (2.95*x*)^2^ – 1.5*x* – 0.5 = 0, so it is approximately equal to 0.341. Thus, 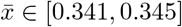, so it is almost constant. In this case the states 1 and 0 are unstable.

**Fig 2.**
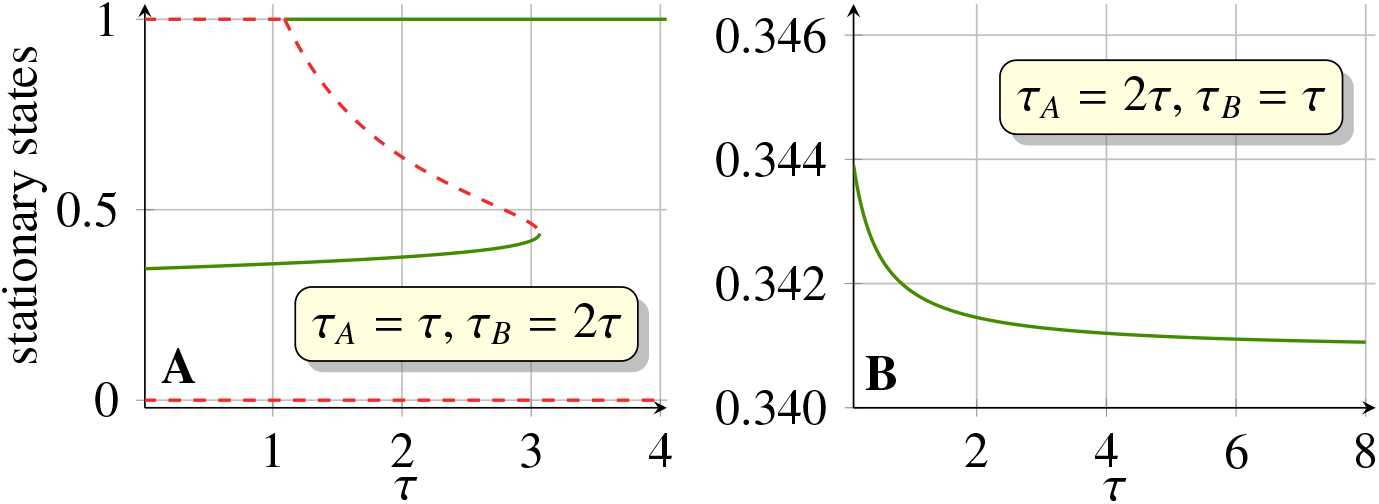
Numerical solutions of *F*(*x*) = 0 for the matrix *U*_2_: stationary state as a function of *τ*, when *τ_A_* = *τ, τ_B_* = 2*τ* (panel A), and when *τ_A_* = *τ, τ_B_* = 2*τ* (panel **B**).

## Conclusion

We studied effects of strategy-dependent time delays on stationary states of evolutionary games.

Recently effects of the duration of interactions between two players on their payoffs and therefore on evolutionary outcomes were discussed by Křivan and Cressman [14]. In their models, the duration of interactions depend on strategies involved. This naturally can be interpreted as strategy-dependent time delays. They showed that interaction times change stationary states of the system.

Another approach is to consider ordinary differential equations with time delays [24]. It was shown that for small time delays, the stationary state is asymptotically stable and at a certain critical time delay, the system undergoes the Hopf bifurcation - the interior state looses stability, oscillations arise. However, it was pointed out in [6] that in the so-called biological model, where it is assumed that the number of players born in a given time is proportional to payoffs received by their parents at a certain moment in the past, the interior state is asymptotically stable for any time delay.

Here we observed a novel behavior in two-player games with strategy-dependent time delays. We showed that interior stationary states depend continuously on time delays. Moreover, at certain time delays, the interior stationary state ceases to exist or there may appear another interior stationary state. Our results are qualitatively similar to those obtained in [14] in a completely different model. We considered only a special case study, more systematic investigations are needed, especially concerning classic games describing social dilemmas. It would be also interesting to analyze strategy-dependent time delays in stochastic dynamics of finite populations.

## Supporting information

### S1 Appendix. Document containing proofs

We provide here propositions, theorems, and their proofs which support results presented in the paper. Additional remarks are also included.

#### Proposition 2.

*If the initial functions φ_x_, φ_p_ are continuous on* [0 – *τ_M_*, 0) *and non-negative, then there exists a unique, non-negative solution of system* (5), (7) *with initial condition* (8) *well defined on the interval* [0, +∞).

*Proof*. The local existence of the solutions follows immediately from a standard theory of delay differential equations, [4]. The non-negativity follows from [1]. To prove the global existence it is enough to use the step method and to observe that on the interval [0, min{ *τ_A_,τ_B_*}] system (5), (7) becomes a system of non-autonomous ordinary differential equations. The equation for *p* becomes linear and can be solved and the equation for *x* then is also linear with respect to *x*(*t*).

Interior stationary states are given by zeros of the function *F*(*x*) defined in (13). Let 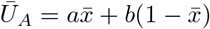 and 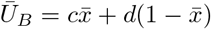.

#### Remark 3.

*We see that for 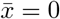 we have*

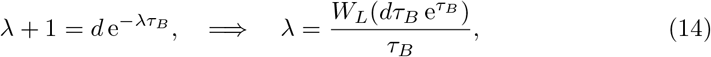

*while for* 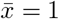 *we have*

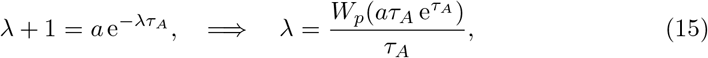

*where W_p_ is the Lambert W function, that is a principle branch of the relation W_p_*(*x*)exp(*W_p_*(*x*)) = *x*.

We show that if *τ_A_* ≠ *τ_B_*, the value of interior stationary state depends on time delays. Moreover, for some pay off matrices and some values of delay, multiple interior stationary states exist. We would like to point out that these relations are not linear and are governed by a non-linear function. Thus, in contrary to the case without time delay or with equal delays (i.e. *τ_A_* = *τ_B_*), adding a constant to a column of pay-off matrix or multiplying the column by a constant changes the value of interior stationary state or even may change the number of interior stationary states. Thus, the intuition from non-delayed games about the asymptotic of dynamics of replicator equation are not necessarily valid for the case of non equal delays.

It turns out, that if the eigenvalue 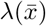 corresponding to the stationary frequency 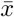 is positive, then the frequency of the given strategy is larger than the frequency of this strategy in a non-delayed case if the delay corresponding this strategy is larger. We have the following

#### Proposition 4.

*Let a < c and d < b. Assume that 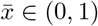 and let* λ *be the leading eigenvalue that corresponds to* 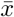. *Then if* λ > 0 *we have*

- *if τ_A_* > *τ_B_ then* 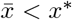;
- *if τ_A_ > τ_B_ then* 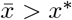;

*Proof*. Assume that *τ_A_* > *τ_B_*. Then the sign of λ is the same as the sign of

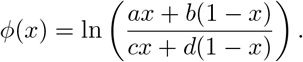

Note, that *ϕ*(*x*\*) = 0 and

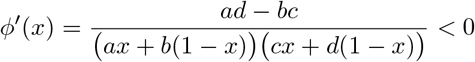

because *ad* < *bc*. Thus, *φ*(*x*) > 0 for *x* < *x*\*, so 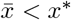. Similarly, if *τ_A_* < *τ_B_* then he sign of λ is opposite to the sign of *ϕ*(*x*) and therefore 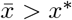.

We will study general properties of *F*, they will help us to determine a number of stationary states of our replicator dynamics. First, we determine conditions that would imply the sign of *F* at *x* = 0 and *x* = 1. Let us define

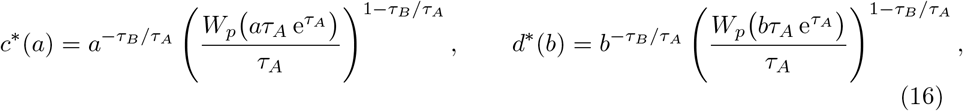

where *W_p_* is the Lambert *W* function, that is a principle branch of the relation *W_p_*(*x*)exp(*W_p_*(*x*)) = *x*.

#### Theorem 5.

*If a < c then F*(1) > 0 *if and only if one of the following condition holds*

i. *τ_A_* > *τ_B_, and a* < 1 *and c* < *c*\*(*a*);
ii. *τ_A_* < *τ_B_, and a* < 1 *or c* > *c*\*(*a*);

*If b > d then F*(0) > 0 *if and only if one of the following condition holds*

i. *τ_A_* > *τ_B_*, *and b* < 1 *or d* < *b*\*(*b*);
ii. *τ_A_* < *τ_B_*, *and b* < 1 *and d* > *b*\*(*b*);

*Proof.* First, we study the sign of *F*(1). It is easy to see that

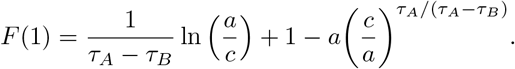

Assume that *τ_A_* > *τ_B_*. Due to the assumption *a* < *c* there exists *z* ∈ (0,1) such that *a* = *zc*. Plugging this into the expression for *F*(1) we obtain

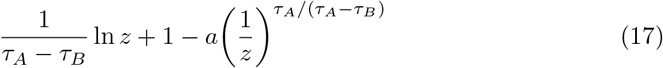

Let us introduce *u* = *z*^*τ_a_*/(*τ_A_* – *τ_B_*)^. Then, the above expression simplifies to

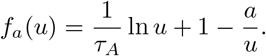

It is easy to see that *f_α_* is a continuous strictly increasing function of *u* and 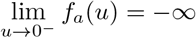. Thus, two possibilities are possible. Either *f_a_*(1) ≤ 0 and *f_a_*(*u*) < 0 for all *u* ∈ (0,1) (thus *F*(1) < 0 for all *a* < *c*) or there exits *u*\* *∈* (0,1) such that *f_α_*(*u*\*) = 0 and *f_a_*(*u*) < 0 for 0 < *u* < *u*\*, and *f_a_*(*u*) > 0 for *u*\* < *u* < 1 (and this implies appropriate sign of *F*(1) depending on the value of a with respect to *c*).

Note, that *f_a_*(1) ≤ 0 is equivalent to *a* ≥ 1. Assume now, that *a* < 1 and we find *u*\*. We have

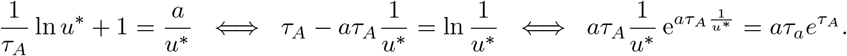

Thus,

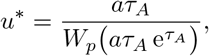

where *W_p_* is Lambert function. Because 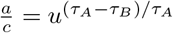, then *F*(1) < 0 if *a* ≥ 1 or *c* > *c*\*(*a*) where *c*\*(*a*) is given by (16). Thus, if *a* > 1 and *c* < *c*\*(*a*), then *F*(1) > 0 and the point (i) is proved. Finally, note that if *τ_A_* < *τ_B_* then in the variable *u* changes from 1 to +∞ instead of from 0 to 1 and very similar arguments leads to the assertion of the point (ii).

Let us calculate the value

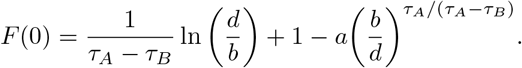

Note that this situation is analogous to the previous one and analogues arguments proves this part of theorem.

If the function *F* is monotonic, then the previous theorem gives us a condition guaranteeing the existence of a solution of *F*(*x*) = 0 on (0,1) that is the existence of an interior stationary state.

#### Theorem 6.

*If a < c, d < b and additionally a <d <c or d<a<b, then the function F is monotonic. Moreover if additionally τ_A_ > τ_B_, then it is decreasing and if τ_A_ < t_B_, then it is increasing.*

*Proof.* It is enough to calculate the first derivative of *F*. It reads

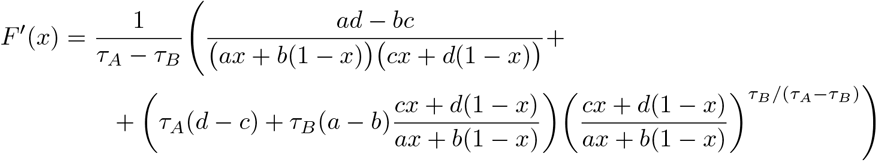

The assumptions guarantee that *ad* – *bc* < 0, *d* – *c* < 0 and *a – b* < 0 and the thesis follows.

#### Corollary 7.

*If a = b or c = d then the function F is monotonic and there is at most one solution of F*(*x*) = 0 *in the interval* [0, 1].

#### Remark 8.

*Theorems 5 and 6 gives a complete description of the existence of the unique stationary frequency 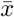 inside the interval* (0,1) *if a<d<c or d < a < b (under the assumption that a < c, d < b)*.

Now, we give a condition that guarantees the existence of the interior stationary point 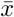 when *τ_B_* is fixed and *τ_A_* converges to zero.

#### Proposition 9.

*Assume that 0 <b < a < c and d = 0. If one of the following conditions*

a. 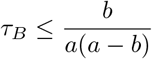 *and a* > 1 *and* 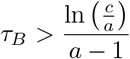
b. 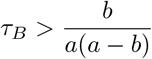 *and b* > 1

*then for τ_A_ < τ_B_ close enough to 0 there exists no interior equilibrium state* 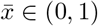.

*Proof*. We check if the equation *F*(*x*) = 0, where *F* is given by (13), has a solution 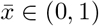 for fixed *τ_B_* and small *τ_A_*. It is easy to see that the function *F* is continuous with respect to *τ_A_* and *x* for *τ_A_* < *τ_B_*. Thus, it is enough to check the existence of solution to *F*(*x*) = 0 for *τ_A_* = 0. In this case, the function *F* simplifies to

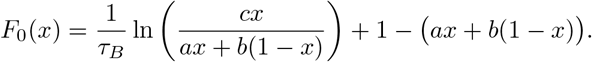

Calculating the derivative of *F*_0_

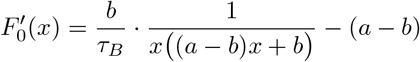

we see that it is a decreasing function of *x* ∈ (0,1) (as *a > b*) and therefore, *F*_0_ is concave. Moreover, 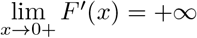 thus

1. if 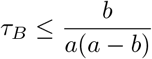 then *F*_0_ is decreasing in (0,1);
2. if 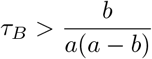 then *F*_0_ has exactly one maximum at (0,1) at the point

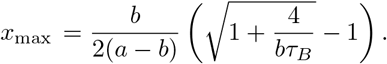

Let us consider the first case. Here, 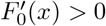 for all *x* ∈ (0, 1), because it is decreasing and 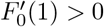. Thus, *F*_0_ is an increasing function of *x* and as lim 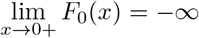 it has a zero in the interval (0, 1) if and only if *F*(1) > 1. An easy calculation allows to derive the condition (a).

Now, let us consider the second case. Some algebraic manipulations leads to

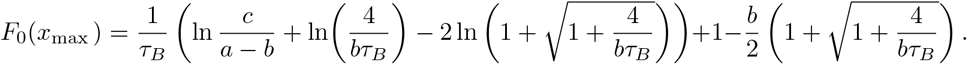

We show that *F*_0_(*x*_max_) approaches to its supreme either for *τ_B_* → +∞ or for 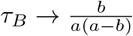. To this end, introduce new variable 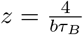 and denote 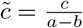. Now, writing *F*_0_ (*x*_max_) with variable *z* we have

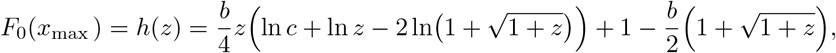

where 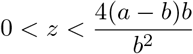 as 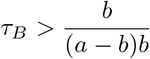. Calculating the derivative of *h* with respect to *z* we obtain

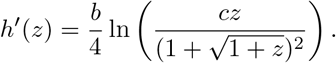

Now it is easy to see that *h*’ has at most one zero for *z* > 0 and it is negative for *z* close to 0. Hence, the function *h* is decreasing for small *z* and it may increase for large *z* having at most one minimum for *z* > 0. Thus, it approaches to its supreme either for *z* → 0 (i.e. *τ_B_* → +∞) or for 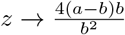. In the latter case, we have

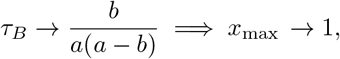

and we arrive at the case considered earlier. On the other hand,

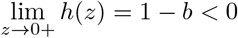

if the condition (b) holds. Thus, *F*_0_(*x*) < 0 for all *x* ∈ (0,1) and no interior equilibrium steady state exists.

#### Remark 10.

*Note, that if* 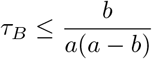 *and either a* < 1 *or* 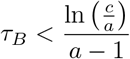 *then there exists exactly one interior equilibrium state* 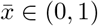 *for τ_A_* < *τ_B_ close enough to 0. On the other hand, if* 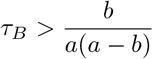 *and b* < 1 *then there exist one or two interior equilibrium states* 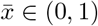 *for τ_A_* < *τ_B_ close enough to* 0 *if τ_B_ is large enough. In fact if a* < 1 *for sufficiently large τ_B_ there exits two equilibrium states* 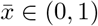.

If *τ_A_* is small enough (for fixed *τ_B_*) or, reversely, if *τ_B_* is large enough (for fixed *τ_A_*), there exists no interior stationary state. Thus, it is possible to calculate these threshold values 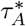 and 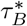. It can be seen that the interior stationary state disappears when it merge with the stationary state 1. Thus, looking for 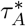 and 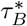 such that *F*(1) = 0, after some algebraic calculation we obtain the implicit formula

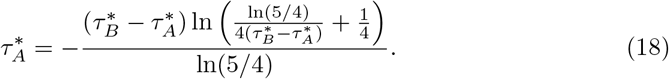

We have that 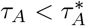 or for 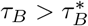 there exists no interior stationary state.

## Acknowledgments

We would like to thank the National Science Centre, Poland, for a financial support under the grant no. 2015/17/B/ST1/00693.

